# Burst c-VEP Based BCI: Optimizing Stimulus Design for Enhanced Classification with Minimal Calibration Data and Improved User Experience

**DOI:** 10.1101/2023.08.18.553900

**Authors:** K. Cabrera Castillos, S. Ladouce, L. Darmet, F. Dehais

## Abstract

The Steady State Visual Evoked Potential (SSVEP) is a widely used technique in Brain-Computer Interface (BCI) research due to its high information transfer rate. However, this method has some limitations, including lengthy calibration time and visual fatigue. Recent studies have explored the use of code-modulated Visual Evoked Potentials (c-VEP) with aperiodic flickering visual stimuli as an alternative approach to address these issues. One advantage of c-VEP is that the training of the model is independent of the number and complexity of targets, which helps reduce calibration time. Nevertheless, the existing designs of c-VEP can be further improved to achieve a higher signal-to-noise ratio, and shorten the selection time and the calibration process. In this study, we propose a novel type of code-VEP called “Burst c-VEP” that consists of brief presentations of aperiodic visual flashes at approximately 3Hz. The rationale behind this design is to leverage the sensitivity of the primary visual cortex to transient changes in low-level stimuli features to reliably elicit distinctive series of visual evoked potentials. In comparison to other types of faster-paced codes, *burst* c-VEP exhibits favorable properties to achieve high decoding performance using convolutional neural networks (CNN). We also explore the attenuation of visual stimuli contrast and intensity to further reduce the perceptual saliency of c-VEP. The proposed solutions were tested through an offline 4-classes c-VEP protocol involving 12 participants. Following a factorial design, participants were asked to focus on c-VEP targets whose pattern (burst and maximum-length sequence) and amplitude (100% or 40% amplitude depth modulation) were manipulated across experimental conditions. Firstly, the full amplitude *burst* c-VEP codes exhibited higher accuracy ranging from 90.5% (with 17.6*s* of calibration data) to 95.6% (with 52.8*s* of calibration data) than its m-sequence counterpart (71.4% to 85.0%). The mean selection time for the two types of codes (1.5*s*) compared favorably compared to existing studies. Secondly, our findings revealed that lowering the intensity of the stimuli barely decreased the accuracy of the *burst* to 94.2% accuracy while yielding a higher subjective visual user experience. Taken together, these results demonstrate the high potential of the proposed *burst* codes to advance BCI beyond the confines of the lab. The collected datasets, along with the proposed CNN architecture implementation, are shared through open-access repositories.

## 1. INTRODUCTION

The Visually Evoked Potentials (VEP) refer to the responses of neuronal populations in the visual cortex to the presentation of visual stimuli. The VEP responses can be reliably captured by surface electroencephalography (EEG). For instance, the presentation of stimuli flickering periodically induces entrainment of the firing rate of neural populations in visual primary cortical areas to the flicker frequency Kritzman et al. (2022); Zhang et al. (2021). This sustained effect has been coined Steady-States Evoked Potentials (SSVEP).

Its rapid onset and robustness, along with the possibility of discriminating a high number of frequencies that can be associated with distinct commands have established the SSVEP as a leading paradigm (reaching unparalleled Information Transfer Rate, ITR) for the design of reactive Brain-Computer Interfaces (BCIs) (Chevallier et al., 2021). To deliver on their promises, several end-user-related issues of SSVEP-based BCI need to be addressed. Firstly, state-of-the-art decoding algorithms used in SSVEP-based BCI rely on template-matching approaches. These template-matching approaches require to be trained on multiple instances of the entire electrophysiological responses associated with each single command (Nakanishi et al., 2018). As such, the higher the number of classes (or “commands”) to be learned by the calibration algorithm, the longer the calibration phase. Considering that the training phase needs to be performed anew for each session, an overly long training time may be a limiting factor to the adoption and retention potentials of SSVEP-based BCI applications. An example of this limitation is illustrated by the study from Chen et al. (2022) which showcases the successful implementation of a 120-commands keyboard SSVEP-based BCI that however requires a 40-minute calibration procedure to be undertaken at the beginning of each session. Secondly, the intensity of flickering visual stimuli has been maximized in an attempt to maximize the EEG responses and henceforth improve classification accuracy (Zemon and Gordon, 2006; Wu and Lakany, 2013; Norcia et al., 2015; Duszyk et al., 2014). Prolonged exposure to these flickering stimuli has reportedly been linked with several inconveniences whose severity ranges from minor visual discomfort (Volosyak et al., 2011) to lasting eye strain (Zhu et al., 2010), from induced mental fatigue to consequential episodes of drowsiness (Cao et al., 2014; Ortner et al., 2011; Patterson Gentile and Aguirre, 2020) and with the additional risk of triggering epileptic seizures in photosensitive individuals (Fisher et al., 2005). Recent studies have demonstrated that SSVEP responses can effectively be elicited by flickers of substantially attenuated contrast to improve user visual comfort without detriment to classification performance (Ladouce et al., 2022, 2021; Lingelbach et al., 2021). Designing repeated visual stimuli that are not intrusive is critical to ensure users’ comfort and safety. Another issue of SSVEP-based BCI lies in the constraints related to its synchronicity requirement. Indeed, decoding which command is selected by the user through template-matching requires the comparison of a definite time series of EEG data spanning from the onset to the offset of the input visual stimulation with a set of trained templates. This approach, therefore, implies knowledge regarding the precise timing at which visual stimuli are not only displayed but also and most importantly being attended to by the user Nagel and Spüler (2019).

Shifting this paradigm to code-VEP (c-VEP) could alleviate several of its flaws. Code-VEP is an emerging paradigm for reactive BCI that substitutes the periodic flickers with aperiodic, binary (zeroes and ones corresponding to black/off and white/on states of the flicker), random code sequences (for a detailed review - see (Martínez-Cagigal et al., 2021)). While the different targets are all derived from the same unique sequence, their phase is shifted. The code sequence is specifically designed to ensure a low correlation between phase-shifted segments. Recent studies have introduced an innovative decoding method that establishes a direct relationship between EEG signals captured on short sliding windows and the elementary bits of c-VEP stimulation, represented as either ‘Black’ (zero) or ‘White’ (one). By sequentially decoding each bit, a complete sequence is generated, which can then be compared to the stimulation patterns of different targets. This approach allows the model to be trained using a limited number of trials, facilitating faster learning. Nagel and Spüler have referred to this approach as “EEG2Code” (Nagel and Spüler, 2019), while Thielen et al. (2021) named it “Re(con)volution”. In a particular scenario, the latter authors demonstrated in their study that the BCI can achieve high accuracy even with minimal or no training data, showcasing the potential for calibration with minimal input.

The bitwise decoding of target stimulation operates on short sliding windows of 0.250ms duration, containing less information compared to template-based approaches that utilize data from a full epoch of stimulation (1s). However, this method necessitates a higher number of short sliding windows. Such a bitwise decoding model requires increased discriminative power and the ability to handle large volumes of data. Convolutional Neural Networks (CNNs) are considered ideal candidates for bitwise decoding of c-VEP, as proposed by (Nagel and Spüler, 2019). CNNs offer numerous advantages in effectively handling the nonlinear characteristics of neural responses and provide exceptional accuracy with remarkably short selection times (typically less than 2 seconds) compared to standard regression approaches (Nagel and Spüler, 2019). However, this approach comes with a cost in terms of CNN computation time that is not compatible with real life scenarios in terms of embedded hardware solutions and duration for the user (Fairclough and Lotte, 2020).

One solution to address this issue is to adapt the design of the c-VEP itself. Currently, the maximum-length sequence (m-sequence) is one of the most popular techniques used for c-VEP implementation (Martínez-Cagigal et al., 2021), which relies on pseudo-random and aperiodic binary sequences alternating plateaus of “ones” (visual stimuli on) and “zeros” (visual stimuli off). By utilizing slow and brief bursts of visual stimuli, one can expect two majors improvements. Firstly, the classification task is simplified into a two-class problem, distinguishing between the presence of the *burst* stimulus (“On”) or its absence (“Off”). This reduction in complexity should reduce computation costs and shorten training times for CNN. Secondly, this approach leverages the sensitivity of primary visual cortex to short-lasting, aperiodic light stimulations to elicit robust event-related responses (Luo and Ding, 2020). While these latter authors overlooked the c-VEP literature and related decoding methods, they took advantage of two continuous sequences of aperiodic flashes placed in foveal and parafoveal vision to tag targets detection with event-related potentials (ERPs).

While c-VEP sequence are deemed as more visually comfortable than SSVEP thanks to their aperiodic nature, this type of visual stimulation still remain salient and straining on the eye in the long run. One solution to improve visual comfort would be to increase the flickering frequency to make the flickers visually transparent to the user. For instance, some c-VEP studies (Martínez-Cagigal et al., 2023; Başaklar et al., 2019) disclosed that increasing the presentation rate of the aperiodic stimuli considerably improve visual user experience. However, these stimuli still remain visible and this approach tend to reach a limit due to reduced orthogonality among templates as they begin to overlap more extensively (Başaklar et al., 2019). An alternative solution to improve user experience and visual comfort is to reduce the contrast and intensity of visual stimuli by lowering their amplitude depth. Stimulus amplitude depth refers to the contrast difference between the two antagonist states of an RVS. In a recent SSVEP study, Ladouce et al. (2022) disclosed that reducing the amplitude depth of VEP considerably improves the user experience while displaying similar high accuracy than full amplitude ones. Adopting a similar approach for c-VEP stimuli could enhance visual comfort since their aperiodic presentation is generally less visually taxing than SSVEP stimuli. This approach has been partially adapted for c-VEP by Gembler et al. (2020), where they introduced the concept of utilizing a quintary m-sequence. Essentially, the quintary m-sequence encompasses five distinct values and employs a range of gray shades for encoding, thereby effectively reducing the intensity of contrast between successive flashes. Their study, along with another recent research by Martínez-Cagigal et al. (2023), revealed improved visual experience while maintaining high accuracy. These results provide evidence supporting the validity of amplitude depth reduction to improve the user experience of VEP-based BCI.

The aim of the present study is to evaluate the effectiveness of a novel *burst* c-VEP sequence design along with amplitude depth reduction in order to improve the user experience of reactive BCI. Building on our previous research on SSVEP (Ladouce et al., 2022, 2021), we opted for a reduction of amplitude depth to only 40% of this amplitude, as previous research demonstrated that this reduction achieves the best compromise between classification accuracy and improved visual comfort. The experiment consisted in a task where participants, equipped with EEG, were required to focus on two different types of c-VEP (burst or m-sequence) for pattern code stimulation. This experiment encompasses two conditions of stimulation: one with 100% amplitude stimulation and the other with 40% amplitude stimulation. By comparing these conditions, we can assess the impact of amplitude reduction on user experience and performance. Our primary objective is to evaluate the efficacy of the *burst* c-VEP compared to the m-sequence c-VEP, while also examining the effects of amplitude reduction. To achieve this, we will measure various performance evaluation metrics, including CNN training time, classification accuracy, and selection time. Additionally, we will investigate how the number of calibration data points affects these metrics. In order to provide a comprehensive analysis, we will also examine evoked responses and inter-trial coherence (ITC) metrics. These analyses can reveal the distinct brain responses elicited by the *burst* c-VEP in comparison to the m-sequence c-VEP. Finally, we conduct subjective assessments to evaluate factors such as visual comfort, visual tiredness, and intrusiveness associated with each code and amplitude. These assessments aim at providing valuable insights into the subjective experiences of participants, complementing the objective performance measures obtained from the experiment.

## 2. MATERIAL AND METHODS

### 2.1. Participants

Twelve healthy volunteers (4 women, mean age: 30.6 years, standard deviation: 7.1), all students and staff at ISAE-SUPAERO with normal or corrected-to-normal vision took part in this study. None of the participants reported any of the exclusion criteria (neurological antecedents and being under psychoactive medication at the time of the study). The study was approved by the ethics committee of the University of Toulouse (CER approval number 2020-334) and was carried in accordance with the declaration of Helsinki. Participants gave informed written consent prior to the experiment. The anonymized data are available at https://zenodo.org/record/8255618.

### 2.2. Stimuli Design

We utilized two types of codes to generate visual stimuli in our study: maximum-length sequences, which are widely recognized as one of the most popular techniques for implementing c-VEP (Martínez-Cagigal et al., 2021), and our newly developed codes called *bursts*.

The *bursts* codes consist of brief flashes (typically less than or equal to 5ms), referred to as *bursts*, with a minimum interval of 200ms between the onset of one *burst* and the onset of the next. Additionally, we incorporated a variable duration between consecutive bursts to create a code rather than a periodic signal. The maximum duration between consecutive flashes is limited to a fixed bound, set at 500 *ms*. More formally, we can represent a *burst* sequence using the compact 4-tuple notation (*f, min, seq, shift*)_*R*_, where *f* is the duration of a *burst, min* is the minimal duration between the onset of two *bursts, seq* = [*t*_1_, …, *t*_*n*_] is the sequence of the variable part between each *burst*, including the minimal duration, *shift* is the phase of the code (circular right-shift), and *R* is the screen refresh rate the code is designed for (as the duration is expressed in frames rather than milliseconds). This notation contains all the necessary information to generate the code data array. Refer to Figure 1 for an illustration of how they relate.

**Figure 1:**
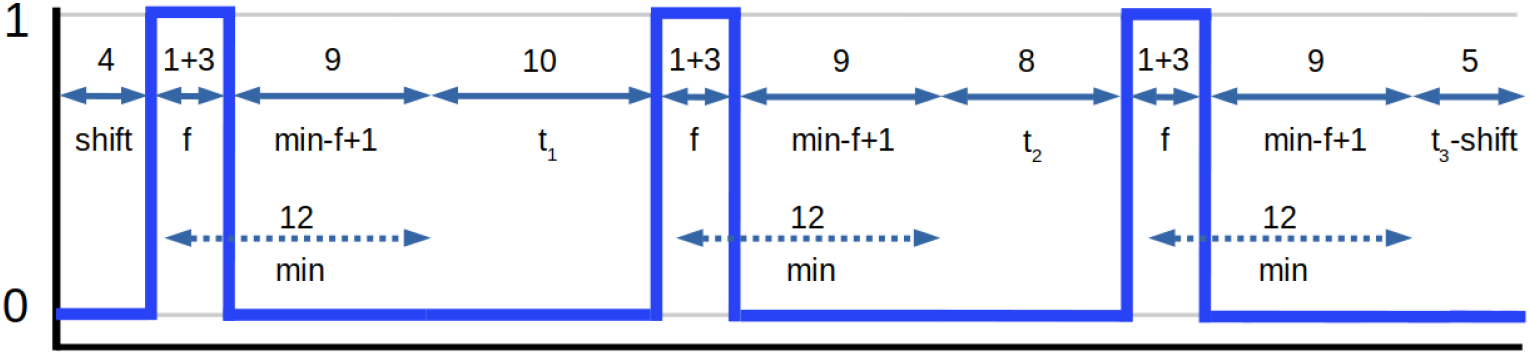
Burst code from compact representation (4, 12, [10, 8, 9], 4)_60_. To create the unfolded sequence, we iterate through all the additional times in the sequence, adding the *burst* frames (*f* = 4), the minimal duration (*min* = 12, minus the burst frames, thus adding only *min* − *f* + 1) and the current additional duration (*t*_1_ = 10, *t*_2_ = 8 and *t*_3_ = 9). The last step is to circularly shift the code to the right, by 4 frames, giving the final *burst* code. Since the total number of frames for this code amounts to 3 × 4 + 3 × 9 + 10 + 8 + 9 = 76 frames and since the code is made for a 60Hz screen refresh rate, the code duration is then meant to be 1.26s

In assessing the correlation across distinct *burst* sequences, we capitalize on the concept of a burst’s “neighborhood” – instances where bursts from divergent codes are in closer proximity, thus exhibiting heightened correlation. By xamining he neighborhood surrounding bursts within one code concerning another, we ascertain their relative closeness. In the context of a code *c*_1_, each individual *burst* undergoes scrutiny for its distance in relation to *bursts* within a separate *c*_2_ code. For this analysis, we establish a defined vicinity around the commencement of a burst (set at 100ms in our instance, as we are looking for clean P100 responses). We then quantify the linear gap between the two bursts, assigning a value of 1 if both bursts initiate exactly simultaneously, and 0 if a burst lies beyond the predetermined neighborhood range. Subsequently, a mean value is computed from the accumulated correlation scores. Should this mean value surpasses a prearranged threshold, it signifies an excessive proximity between the two codes in their present phase, prompting their classification as overly convergent.

One may raise the question of potential duplication in counting a burst within *c*_2_, should it find itself in the vicinity of two *c*_1_ bursts. This scenario, however, does not impart symmetry to the correlation under consideration. Our specific circumstances preclude such an occurrence: with an inter-burst interval of 200ms and a neighborhood zone spanning 100ms, even under the most unfavorable conditions (wherein no temporal gap exists between the two bursts), there remains an absence of overlap within the adjoining zones of consecutive bursts within the same code. Thus, it becomes evident that a burst within *c*_2_ cannot simultaneously inhabit the neighboring zones of both bursts, thereby confirming the adequacy of this straightforward correlation definition in our context.

With our correlation function in place, we embark on the generation of our *burst* codes, initiating a systematic assessment of correlations through phased adjustments. Within each code pair, our focus narrows to the paramount correlation score derived from the array of scores originating from these shifted codes. A pivotal facet, this peak correlation score, is then juxtaposed against an established threshold. When the resulting score falls below this predefined threshold, it signifies a diminished likelihood of misinterpretation, an outcome resilient against variations in relative phases. As a result, the crux of our code selection process revolves around upholding minimal pairwise correlation—a fundamental criterion guiding our final code choices.

For our experiment we thus use the following *burst* codes, with Figure 2 showing their visual representation:

**Figure 2:**
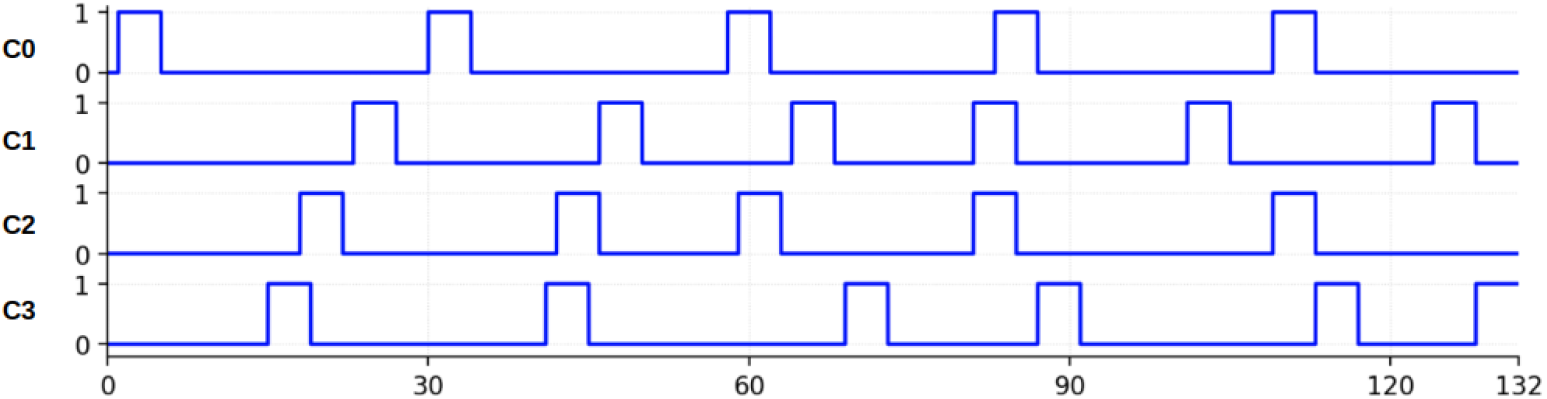
Illustration depicting our four experimental *burst* codes: *C*1, *C*2, *C*3, and *C*4. Each code consists of 132 frames, equivalent to a duration of 2.2 seconds, designed for optimal display on a 60Hz screen, as required for our experiment.

- C1 = (4, 12, [16, 15, 12, 13, 11], 1)_60_
- C2 = (4, 12, [10, 5, 4, 7, 10, 18], 23)_60_
- C3 = (4, 12, [11, 4, 9, 15, 28], 18)_60_
- C4 = (4, 12, [13, 15, 5, 13, 2, 6], 15)_60_

Concerning the m-sequences, we drew upon codes originally formulated in a preceding study, achieving a 98% accuracy within an 11-class problem (Darmet et al., 2023). Our approach involved the utilization of a Fibonacci-type linear feedback shift register (LFSR), employing two distinct polynomials: *x*9 + 1 and *x*9 + *x*6 + *x*3 + 1. The initial states chosen were 11111110011, 01111010011, 10110110011, 10111110011, and 01111110011. This configuration enabled the construction of a codebook containing m-sequence-like codes. The codes derived from our codebook underwent segmentation into subsequences, each spanning 132 frames. Given the four-class nature of our experiment, we ultimately used the initial four codes as the most optimal candidates, after making sure that the correlation between them and across all their phased versions is not higher than 0.5.

Finally, in addition to the variation of the type of code, we presented both *m-sequences* and *burst* c-VEP at two different levels of amplitude depth of stimulation: 100% amplitude depth (i.e., maximal screen luminosity) and 40% amplitude depth (see Figure 3 - right). Finally, in addition to the variation of the type of code, we presented both *m-sequences* and *burst* c-VEP at two different levels of amplitude depth of stimulation: 100% amplitude depth (i.e., maximal screen luminosity) and 40% amplitude depth (see Figure 3 - right). Since the background was set to a medium gray (50% luminosity), the 40% amplitude is between this medium gray and full screen luminosity.

**Figure 3:**
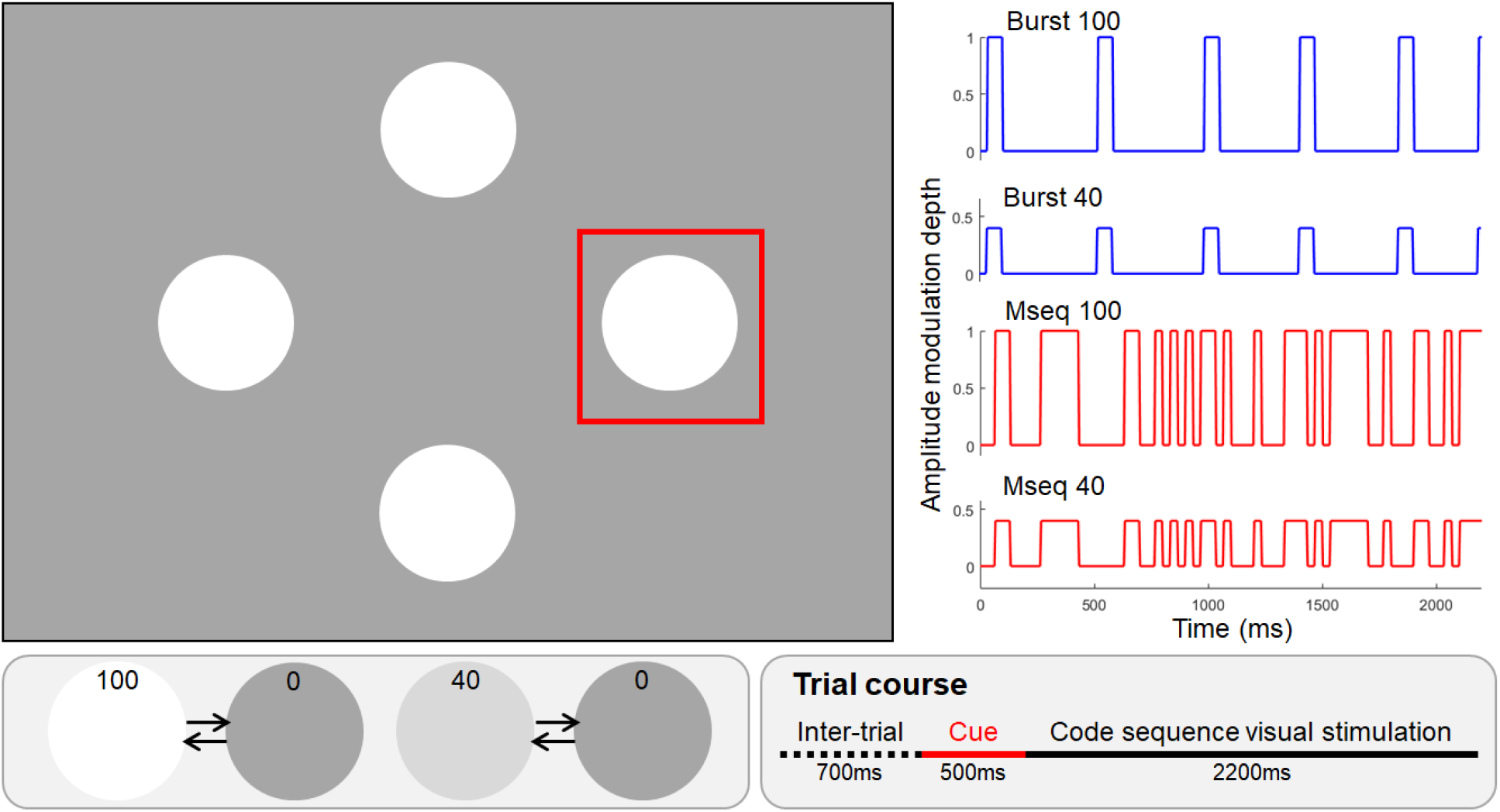
Left: the presentation of *burst* and m-sequence based c-VEP varied in amplitude depth, with options for either maximum (100%) or reduced to 40%, aiming to enhance visual comfort. Top-right (blue): the brief aperiodic *burst* based c-VEP presented at a rate of approximately 3Hz. Bottom-right (red): The m-sequence is an aperiodic binary sequence that alternates short-lasting plateaus of visual stimuli “on” and “off” at a frequency of approximately 10Hz. The x-axis is time (in milliseconds). Bottom-left: Representation of the two alternating states of visual stimuli for both amplitude modulation depth (100 and 40%). Bottom-right: time course of a trial, an inter-trial interval of 700ms separates the current trial from the previous one. A red-bordered square cue then appears for 500ms around the stimulus to be attended. The cue is followed by the onset of visual stimulation where all the different codes corresponding to the current conditions (Burst100, Burst40, Mseq100, Mseq40) are presented for 2200ms.

### 2.3. Experimental Protocol

Participants were comfortably sat and requested to read and sign the informed consent. Once equipped with the EEG system, volunteers were asked to focus on four targets that were cued sequentially in a random order for 0.5s, followed by a 2.2s stimulation phase, before a 0.7s inter-trial period. The cue sequence for each trial was pseudo-random and different for each block. After each block, a pause was observed and subjects had to press the space bar to continue. In total, each volunteer was presented fifteen four-trial blocks for each of the four conditions (burst or m-sequence × 40% or 100%) see Figure 3 - left. The task was implemented in Python using the Psychopy toolbox ^1^. The four flickers were all 150 pixels wide discs, without borders, and were presented on the following LCD monitor: Dell P2419HC, 1920 × 1080 pixels, 265*cd*/*m*2, and 60Hz refresh rate. After completing the experiment and removing the EEG equipment, the participants were asked to provide subjective ratings for the different stimuli conditions. These stimuli included *burst* c-VEP with 100% amplitude, *burst* c-VEP with 40% amplitude, m-sequences with 100% amplitude, and m-sequences with 100% amplitude. Each stimulus was presented three times in a pseudo-random order. Following the presentation of each stimulus, participants were presented with three 11-points scales and were asked to rate the visual comfort, visual tiredness, and intrusiveness using a mouse. In total, participants completed 12 ratings (3 repetitions x 4 types of stimuli) for each of the three scales.

### 2.4. EEG Pre-processing

EEG data were recorded using a BrainProduct LiveAmp 32 active electrodes wet-EEG setup with a refresh rate of 500Hz to record the surface brain activity. The 32 electrodes followed the 10-20 international system. The ground electrode was placed at the FPz electrode location and all electrodes were referenced to the FCz electrode. The electrode impedances were brought below 25*k*Ω before the recording. EEG data and markers were synchronized during recording using Lab Streaming Layer ^2^. EEG analyses were conducted only on a subset of specific electrodes: 01, 02, 0z, Pz, P3, P4, P8, and P9. The raw continuous EEG data were average re-referenced. An IIR cut-band filter between 49.9 and 50.1Hz of order 16 was then applied to remove line noise. The raw continuous data were then epoched from 0 to 2.2s around timestamps of flickers onset. Finally, a baseline removal was performed on the epochs to remove eventual slow drifts.

### 2.5. Convolutional Neural Network and Pattern Decoding

To decode the brain response on EEG data, we leverage the power of Convolutional Neural Network (CNN) such as EEG2Code (Nagel and Spüler, 2019). The CNN architecture^3^ used in this study is the same as in our previous work (Darmet et al., 2023), which exhibited better performance in terms of selection time and accuracy compared to the original EEG2Code (Nagel and Spüler, 2019).

To predict the target the subject intends to select, the decoding process consists of two phases. Firstly, short windows of EEG data are input to the CNN to decode the code pattern. Secondly, this decoded pattern is compared to the stimulation patterns of the targets using Pearson correlation. The CNN operates on a 250 ms sliding window with 2 ms steps, adhering to the EEG sampling rate of 500 Hz. However, the actual state switch rate of the simulation is slower due to the screen refresh rate limitation of 60 Hz. Thus, downsampling of the decoded pattern to match the screen refresh rate is performed using majority voting.

In the second phase, when the decoded pattern reaches a minimal length of 0.7 s, a Pearson correlation coefficient is computed between the downsampled binarized decoded pattern and the different target templates. The label of the highest correlating template is temporarily considered as the prediction. The decoded pattern is then expanded with the next stimulation decoding using the following EEG data (i.e., the downsampled next 8 windows), and a new temporary prediction is made. If the system consecutively outputs the same temporary prediction 60 times, a closing prediction for the trial is triggered. Computation is stopped, and the classification starts for the next trial. In cases where no closing prediction could be made after 2 s, the system outputs -1, indicating no prediction, and therefore does not issue any command.

We evaluated the number of needed calibration data with an iterative approach: starting with one block of calibration (2.2*s* × 4 stimuli × 1 block = 8.8*s* of training data) and gradually increasing it to six blocks of calibration (2.2*s* × 4 × 6 = 52.8*s* of training data). This approach allowed us to compare the effects of two stimuli (burst versus m-sequence) and amplitude depth reduction on CNN training time, 4-class selection time, and 4-class decoding accuracy. To simulate an online setting, we performed sequential train/test splits in each of these computations. The epochs were segmented into windows of 250ms with a stride of 2ms (one data point of EEG data sampled at 500Hz). We computed the standard deviation within each time window of the training dataset enabling standard deviation normalization. This normalization value was then applied to the testing data as well. Additionally, we allocated 10% of the calibration data for validation purposes. This subset was used to monitor the loss and accuracy during the CNN training process.

### 2.6. VEP Analysis

The effect of experimental manipulations on Visual Evoked Potentials (VEP) responses was investigated to better understand the neural underpinnings that may explain differences in terms of classification performance. For this purpose, the continuous EEG data was bandpass filtered between 0.1 and 40Hz (FIR, 16501 filter order, cutoff frequencies at 0.05 and 40.05Hz) and then average rereferenced. The onset of singular visual stimuli (within the whole sequence) was defined based on the timings at which the code sequences alternated from the dark to the bright states. The continuous EEG data were then segmented into epochs around the onset of visual stimulation onset (from -0.5 to 1s). The VEP responses were averaged across trials for each electrode and participant. The mean amplitude of the VEP (computed over a 60 to 110 time window) and its peak latency were extracted from the resulting waveforms at electrode Oz over which the VEP response is most prominent.

## 3. RESULTS

Statistical analyses were carried out with JASP 0.16 software ^4^. Repeated measures ANOVA were computed for each subjective and objective metric with types of codes and luminance as within factor. The Greenhouse-Geisser correction was applied when the assumption of sphericity was violated. The Bonferroni test was used for all post-hoc comparisons. Significance level was set at *p* < 0.05 for all analyses.

### 3.1. Subjective Results

We ran a series of repeated measures ANOVA to assess the effect of types of codes and amplitude depth on our subjective metrics (visual comfort, mental tiredness and intrusiveness) when considering 6 blocks of training data (4 flickers × 2.2*s* × 6 blocks = 52.8*s*). Figure 4 shows a visualization of the subjective results.

**Figure 4:**
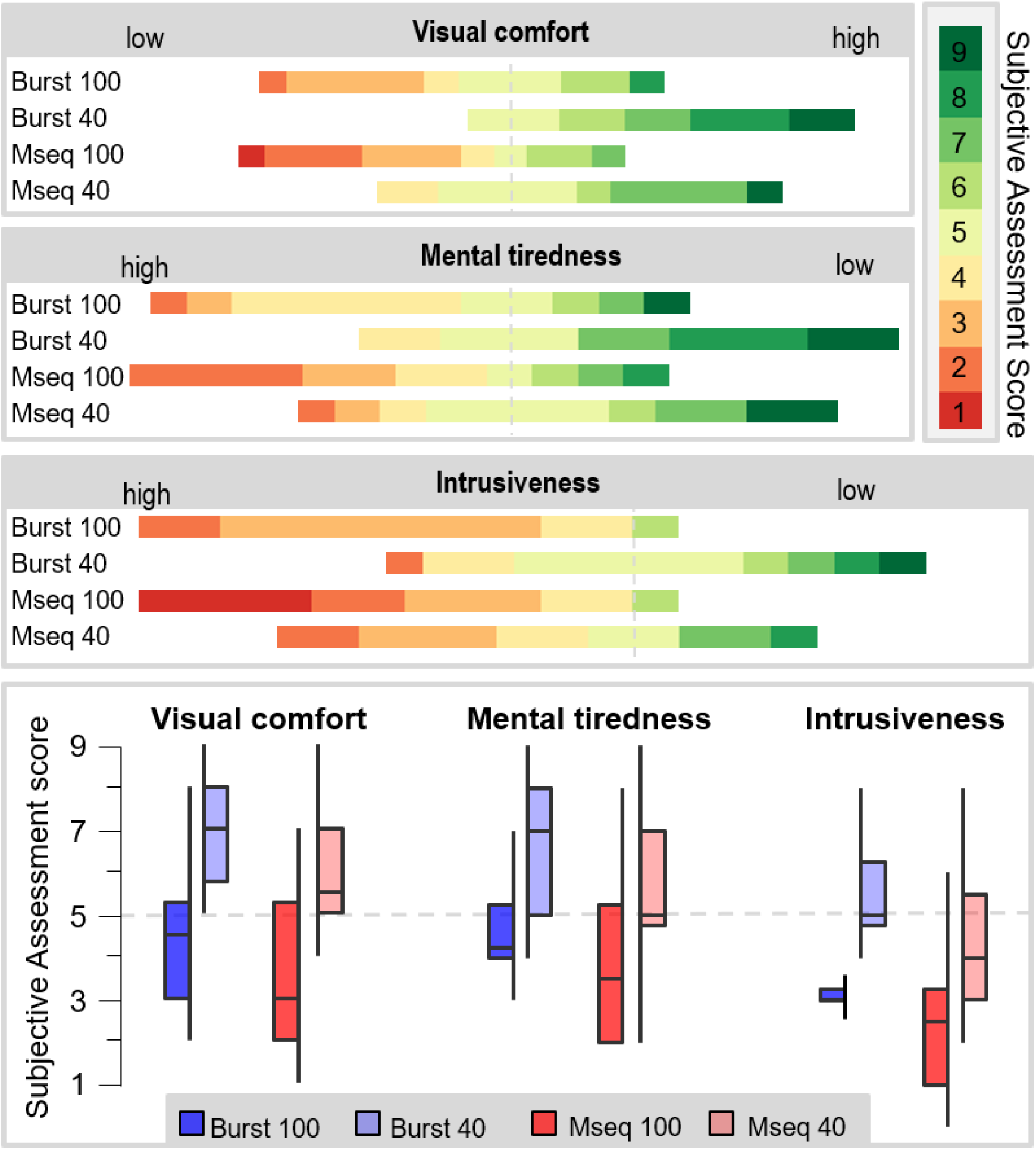
Distribution (N = 12) of subjective assessment of the different visual stimulation code sequences (*Burst 100, Burst 40, Mseq 100, Mseq 40*) across three dimensions: visual comfort, mental tiredness, and presence. All code sequences were rated several times by the participants and the rounded average of scores is presented discretely on the color-coded bar plots (ranging from red for the lowest, to yellow for neutral, to green for the highest ratings). The proportion of individual scores attributed to an item is reflected by the length of the colored bars. For comparison purposes, all score distributions are centered on the neutral rating (5 on the 9-points scale) which is denoted by the dotted lines.

#### 3.1.1. Visual Comfort

A 2×2 repeated measure ANOVA, with factors of c-VEP type (burst, m-sequence) and amplitude depth (low, high), disclosed a main effect of the type of code on visual comfort (F(1,11)= 7.18, *p* = 0.021, 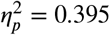) and a main effect of the amplitude depth on visual comfort (F(1,11)= 28.03, *p* = 0.001, 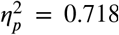) but no significant interactions (F(1,11)= 0.407, *p* = 0.53, 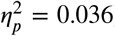. Post-hoc analyses revealed that the *burst* c-VEP led to better visual comfort than their m-sequence counte^*p*^rpart (*p* = 0.021, Cohen’s d = 0.522). Post-hoc analyses also revealed that lowering the amplitude depth led to increased visual comfort (*p ≤* 0.001, Cohen’s d = 1.418).

#### 3.1.2. Mental Tiredness

A 2×2 repeated measure ANOVA, with factors of c-VEP type (burst, m-sequence) and amplitude depth (low, high), highlighted a main effect of the type of code on mental tiredness (F(1,11)= 10.043, *p* = 0.009, 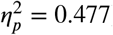) and a main effect of the amplitude depth on mental tiredness (F(1,11)= 16.584, *p* = 0.002, 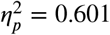) but no significant interactions (F(1,11)= 0.340, *p* = 0.571, 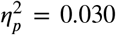). Post-hoc analyses revealed that the *burst* c-VEP led to lower mental tiredness than the m-sequence c-VEP^*p*^ (*p* = 0.009, Cohen’s d = 0.437) and that reducing the amplitude decreased mental tiredness (*p* = 0.002, Cohen’s d = 0.854).

#### 3.1.3. Intrusiveness

A 2×2 repeated measure ANOVA, with factors of c-VEP type (burst, m-sequence) and amplitude depth (low, high), revealed a main effect of the type of code on intrusiveness (F(1,11)= 7.18, *p* = 0.02, 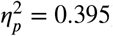) and a main effect of the amplitude depth on intrusiveness (F(1,11)= 31.93, *p ≤* 0.001, 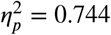) but no significant interactions (F(1,11)= 0.256, *p* = 0.62, 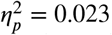). Post-hoc analyses revealed that the *burst* c-VEP were less intrusive than the m-sequence c-VEP (*p* = 0 02, Cohen’s d = 0.515) and that reducing the amplitude also led to decreased intrusiveness (*p* ≤ 0 001, Cohen’s d = 1.201).

### 3.2. BCI Performances

We ran a series of repeated measures ANOVA to assess the effect of types of codes and amplitude depth on our performance metrics (classification accuracy, selection time, CNN model training time, Information Transfer Rate - ITR) when considering 6 blocks of training data (4 flickers × 2.2*s* × 6 blocks = 52.8*s*). We also ran some descriptive analyses to report the effect of the number of blocks of training trials (from one block to 6 blocks) on classification accuracy and CNN training time.

#### 3.2.1. Classification Accuracy

A 2×2 repeated measures ANOVA, with factors of c-VEP type (burst, m-sequence) and amplitude depth (low, high), revealed a significant main effect of the type of code on classification accuracy (F(1,11) = 16.62, *p* = 0.002, 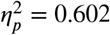) and a significant main effect of the amplitude depth on classification accuracy (F(1,11) = 9.86, *p* = 0.009, 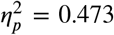). However, no significant interaction of these variables was observed on classification accuracy (F(1,11) = 2.49, *p* = 0.14, 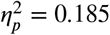). Post-hoc analyses indicated that the *burst* c-VEP resulted in higher accuracy compared to their *m-sequence* counterparts (*p* = 0.002, Cohen’s *d* = 1.177). Similarly, post-hoc analyses revealed that reducing the amplitude depth led to a decrease in classification accuracy (*p* = 0.009, Cohen’s d = 0.907) - see fig. 5 (upper left).

**Figure 5:**
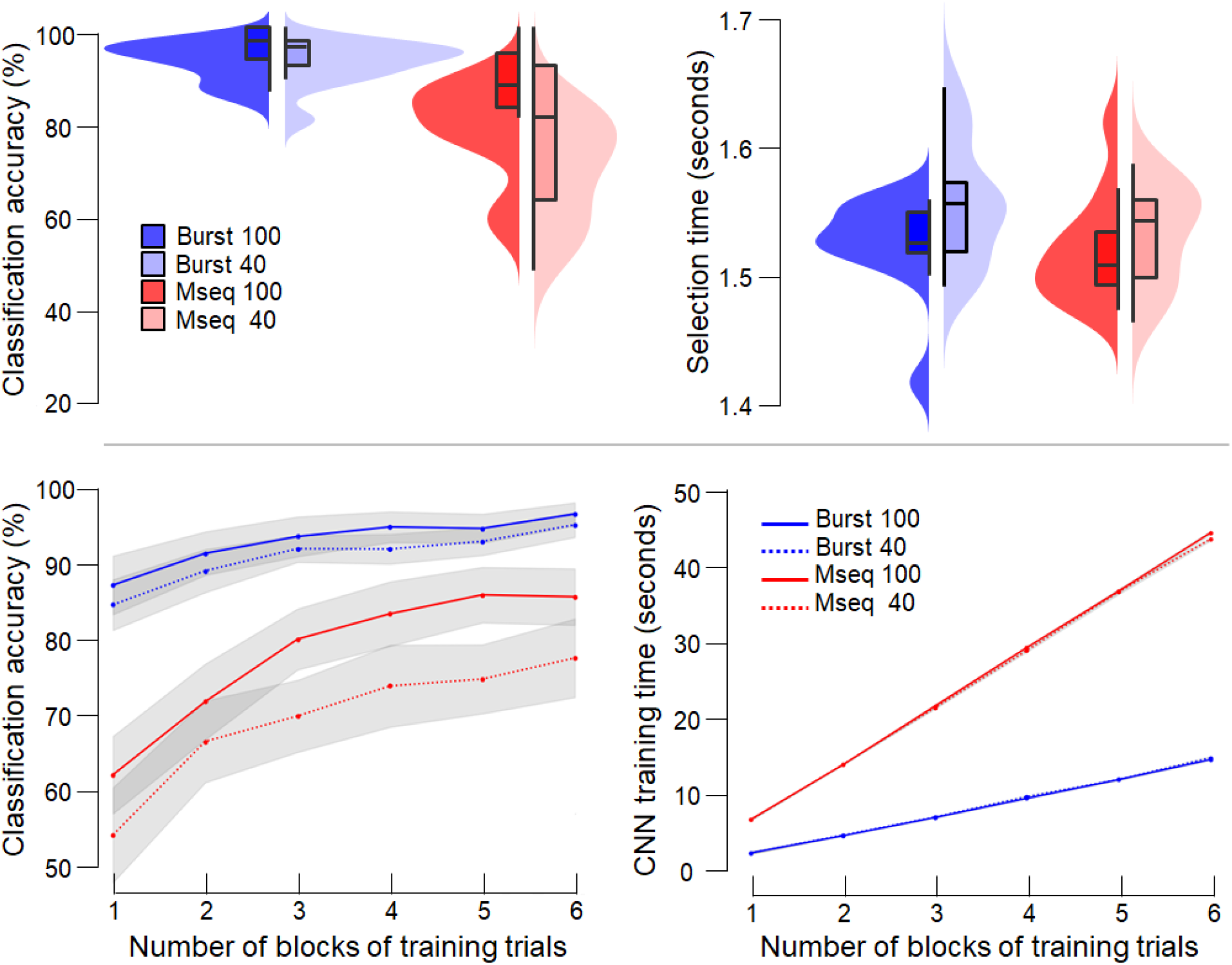
Performance assessment of BCI on the 4-class problem across code sequences (Burst 100, Burst 40, Mseq 100 and Mseq 40) through different metrics. Top-left: Classification accuracy with 6 trials of training data across visual stimulation. Top-right: Comparison of output commands selection time achieved by the CNN with 6 trials of training data per class. Bottom-left: Classification accuracy achieved by the CNN as a function of the number of training trials per class for each code sequence. Bottom-right: CNN training time as a function of the number of training trials. In the present experiment, each additional training trial collected lengthened the calibration time by 8.8 seconds (4 classes x (2.2s stimuli duration)).

**Figure 6:**
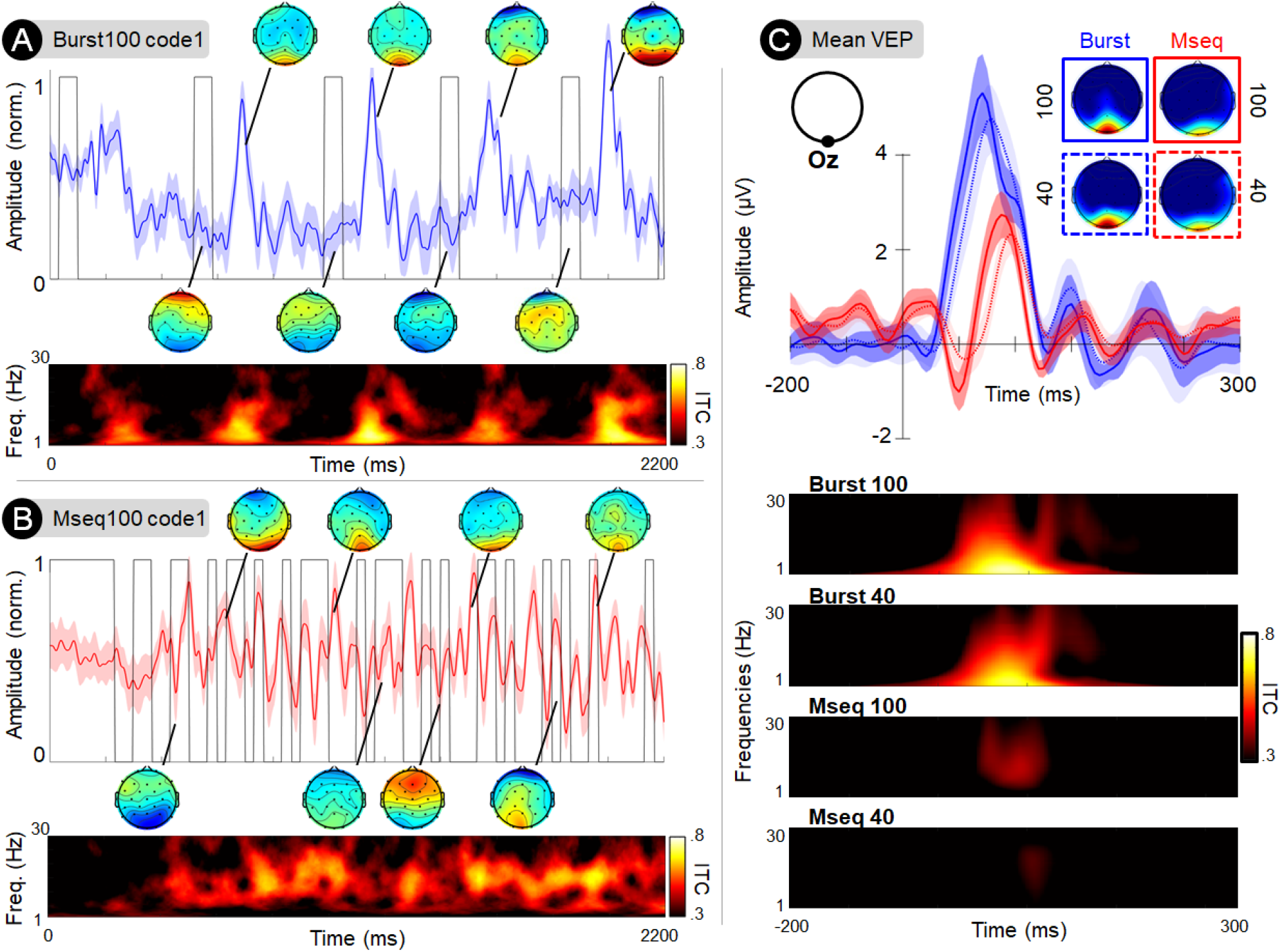
Grand average (N=12) EEG responses recorded at occipital electrode Oz elicited by the presentation *burst* and m-sequence code sequences. A) The blue line presents the grand average Visually Evoked Potentials (VEP) response to the first *burst* code-VEP sequence (black line) lasting for 2.2 seconds. The VEP amplitude was normalized and the shaded area represents the VEP amplitude standard error across individuals computed over time samples. The scalp maps present the instantaneous topographical distribution of the VEP responses at different phases of the stimuli input function. The bottom subplot presents the Inter-Trial Coherence (ITC) of VEPs computed across frequencies (y-axis) and over time (x-axis), where brighter shades denote higher ITC. B) Similarly to subplot A, the red line presents the grand average VEP response to the first m-sequence code-VEP (black line) lasting for 2.2 seconds. C) Top: The line plots present the grand average VEP responses locked to the onset of visual stimulation for both types of code sequences (burst in blue and m-sequence in red) and amplitude modulation depth (100% amplitude in solid line and lower and 40% amplitude in dotted line). The scalp maps present the spatial distribution of average VEP response over the VEP time window (150 to 210ms after stimulus onset). The bottom plots highlight the ITC across all four experimental conditions.

#### 3.2.2. Selection Time

A 2×2 repeated measures ANOVA, with factors of c-VEP type (burst, m-sequence) and amplitude depth (low, high), did not reveal a significant main effect of the type of code on selection time (F(1,11) = 1.29, *p ≤* 0.028, 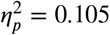) or a significant main effect of the amplitude depth on selection time (F(1,11) = 4.26, *p* = 0.06, 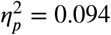). Additionally, no significant interaction was observed on selection time (F(1,11) = 0.66, *p* = 0.43, 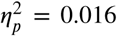 - see fig. 5 (upper right)).

#### 3.2.3. CNN Training Time

A 2×2 repeated measures ANOVA, with factors of c-VEP type (burst, m-sequence) and amplitude depth (low, high), revealed a significant main effect of the type of code on CNN training time (F(1,11) = 36472.86, *p* < 0.001, 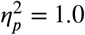). However, no main effect of the amplitude depth on CNN training time was observed (F(1,11) = 15804.50, *p* < 0.001, 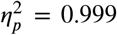). Additionally, no significant interactions were found on CNN training time (F(1,11) = 0.83, *p* = 0.04, 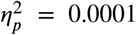). Furthermore, a significant interaction was observed on selection time (F(1,11) = 7.92, *p* = 0.01, 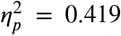). Post-hoc analyses indicated that the *burst* c-VEP with full amplitude and 40% amplitude resulted in sh^*p*^orter training time compared to their *m-sequence* counterparts (*p* < 0.001). Additionally, the full amplitude *m-sequence* exhibited shorter training time compared to the 40% amplitude *m-sequence* (*p* = 0.03).

#### 3.2.4. Information Transfer Rate

A 2 × 2 repeated measures ANOVA, with factors of c-VEP type (burst, m-sequence) and amplitude depth (low, high), revealed a significant main effect of the type of code on Information Transfer Rate (ITR) (F(1,11) = 30.144, *p* < 0.001, 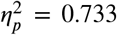) and a significant main effect of the amplitude depth on ITR (F(1,11) = 8.892, *p* = 0.012, 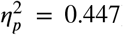). However, no significant interactions were found on ITR (F(1,11) = 0.529, *p* = 0.482, 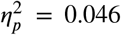). Post-hoc analyses revealed that the *burst* c-VEP resulted in higher ITR compared to their *m-sequence* c^*p*^ounterparts (*p* < 0.001, Cohen’s d = 1.585). Similarly, post-hoc analyses revealed that reducing the amplitude depth led to a decrease in ITR (*p* = 0.012, Cohen’s d = 0.861). The mean ITR values for each condition were as follows: *burst* 100 (mean = 62.713, SD = 11.58), *burst* 40 (mean = 67.49, SD = 11.75), *m-sequence* 100% (mean = 38.83, SD = 23.9),*m-sequence* 40% (mean = 48.7, SD = 19.08).

#### 3.2.5. Impact of Training Trial Blocks on Classification Performance

The results we reported in the previous section on the effects of code type and amplitude were all given with 6 calibration blocks as training data. We further explored the impact of the amount of training data fed to the model for its training. We thus incrementally reduced the number of calibration blocks dedicated to training, from 6 (equivalent to 52.8 seconds) to 1 (equivalent to 8.8 seconds of data), the other being used for testing. The descriptive results of our analysis reveal compelling patterns. Notably, the accuracy of the *burst* codes consistently maintains a level above 85%, surpassing the accuracy achieved by the m-sequence codes. This observation is visually represented in Figure 5, located in the lower left quadrant. Moreover, in terms of training time, the *burst* codes demonstrate a significant advantage over the m-sequence codes. The CNN training time associated with the *burst* codes is consistently at least four times faster than that of the m-sequence codes. This disparity is graphically depicted in Figure 5, displayed in the lower right quadrant. Our descriptive findings suggest that the utilization of *burst* codes not only ensures robust classification accuracy, consistently surpassing 85%, but also offers a substantial reduction in CNN training time compared to the usage of m-sequence codes.

### 3.3. Visual Evoked Potentials Results

We ran a series of repeated measures ANOVA to assess the effect of types of codes and amplitude depth on our the visual evoked potentials metrics (amplitude, latency and inter-trial coherence) when considering 6 blocks of training data (4 flickers × 2.2*s* × 6 blocks = 52.8*s*).

#### 3.3.1. Amplitude

A 2 × 2 repeated measure ANOVA, with factors of c-VEP type (burst, m-sequence) and amplitude depth (low, high), revealed a main effect of code sequence type on VEP amplitude (F(1,11)= 28.25, *p ≤* 0.001, 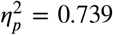) but no main effect of the amplitude depth (F(1,11)= 2.616, *p* = 0.123, 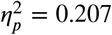) and no interaction between these two factors (F(1,11)= 0.004, *p* = 0.949, 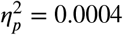). The *burst* code sequences evoked significantly larger VEPs than the m-sequence code sequences (*p ≤* 0.0001, Cohen’s d = 1.637).

#### 3.3.2. Latency

A 2 × 2 repeated measure ANOVA, with factors of c-VEP type (burst, m-sequence) and amplitude depth (low, high), revealed a main effect of code sequence type on VEP latency (F(1,11)= 46.485, *p* <= 0.001, 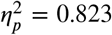) and a main effect of amplitude depth (F(1,11)= 7.888, *p* = 0.019, 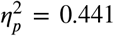). There was no interaction effect on VEP latency between these two factors (F(1,11)= 4.047, *p* = 0.072, 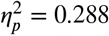). The latency of VEPs responses was found to be significantly earlier in response to *burst* than m-sequence^*p*^ codes (*p ≤* 0.001, Cohen’s d = 1.794). Moreover, VEP responses appeared earlier following the onset of full amplitude visual stimulation than in the case of reduced amplitude stimuli (t(11)=2.808, *p* = 0.019, Cohen’s d = 0.988).

#### 3.3.3. Inter-Trial Coherence

A 2 × 2 repeated measure ANOVA, with factors of c-VEP type (burst, m-sequence) and amplitude depth (low, high), revealed a main effect of code sequence type on theta-alpha ITC (F(1,11)= 62.036, *p* <= 0.001, 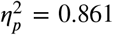) and a main effect of amplitude depth (F(1,11)= 6.752, *p* < 0.05, 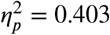). There was an interaction effect on VEP ITC between these two factors (F(1,11)= 12.030, *p* < 0.01, 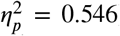). Post-hoc analyses revealed that the *burst* c-VEP induced higher ITC than their m-sequence counterpart (*p* < 0. 001, Cohen’s d = 1.790). The ITC coefficient of VEPs responses was higher in response to full amplitude visual stimulation than in the case of reduced amplitude stimuli (*p* = 0.027, Cohen’s d = 0.288). There was no significant difference in ITC between the burst 100 and 40 conditions (*p* = 1.00, Cohen’s d = 0.094). In contrast, a significant decrease in ITC was observed when the amplitude depth of m-sequence codes was decreased from 100 to 40% (*p* = 0.009, Cohen’s d = 0.481).

## 4. Discussion

The objective of this study was to evaluate the effectiveness of a new type of c-VEP pattern called *burst*. These code sequences consist of short-lasting flashes presented aperiodically at a frequency of approximately 3Hz. Additionally, to enhance the comfort level associated with the presentation of these codes, we proposed to reduce their amplitude depth. In light of this, we designed an experiment to compare and benchmark these codes against the more conventional m-sequence c-VEP. Furthermore, we manipulated two levels of amplitude depth (100% or 40%) to investigate their impact on BCI performance and user comfort.

In order to compare the two types of codes (m-sequence and burst), let’s first consider conditions in which stimuli were presented at maximal amplitude depth. Our findings indicated that the *burst* c-VEP outperformed the m-sequence code on most of the classification performance metrics. When using 6 blocks of training data (amounting to 52.8 seconds of calibration data), the full amplitude *burst* c-VEP exhibits a significantly higher classification accuracy (95.6%) compared to the m-sequence codes (85.0%). Moreover, the proposed *burst* codes demonstrated greater robustness to reductions of training data compared to the m-sequence. With no more than two blocks of calibration (corresponding to only 17.2*s* of training data acquisition), the classification accuracy of neural responses to *burst* c-VEP sequences remained high, reaching 90.5%, while classification accuracy related to the m-sequence codes dropped substantially to 71.4%. This finding highlights the relevance of the *burst* code design in reducing calibration time without sacrificing classification accuracy. To provide context, the original EEG2Code study, which first utilized CNN to decode the m-sequence, required 384 *s* of calibration data for training (Nagel and Spüler, 2019). Similarly, a previous study conducted by our research group (Darmet et al., 2023) using a similar m-sequence stimulus design and CNN architecture as in the present research required 169.4 *s* of calibration data. In a recent comparison, slightly slower-paced aperiodic code sequences were shown to outperform regular m-sequences c-VEP in terms of accuracy and calibration time, reducing the latter down to a range of 1 to 3 minutes (Gembler et al., 2020; Verbaarschot et al., 2021). Furthermore, a recent study Thielen et al. (2021) showcased that c-VEP-based BCI can be calibrated with just a few trials at the start of the session, or even without any training data altogether. This intriguing approach has the potential to be incorporated into the decoding approach used in this work, further reducing calibration time and, as such, improving the overall user experience. On a related issue, the training time for the CNN monotonically increases as a function of training data. However, the steepness of the slope characterizing this relationship is significantly higher for the m-sequence than the *burst* amounting to 40 seconds against 15 when using 6 blocks of training trials. Both of these measures clearly demonstrate that the *burst* stimulus design substantially reduces calibration and CNN training time. This is an important consideration within the frame of non-invasive BCI development as a new calibration phase is required at the start of each session. Therefore, reducing the initialization phase of the BCI setup is critical to ameliorate user experience and ensure user retention.

Another critical aspect of BCI performance which is closely tied to user experience is how long it takes for the system to identify the command selected by the user and perform the associated action. In this study, the mean selection decoding time of the two types of codes was similar (1.5*s*). This fast decoding time took advantage of the CNN approach both in terms of reactivity and flexibility. The proposed CNN implementation outperformed the EEG2code study that used CNN (1.7*s*) (Nagel and Spüler, 2019), and showed faster decoding times than the study by Dehais et al. (2022) which reported selection duration varying between 1.6*s* to 2.6*s* depending on the workload condition. It should also be noted that selection times achieved through CNN classification approach are significantly shorter than those achieved with the canonical correlation analysis approach, where the best performances reported range between 3.1*s* and 3.8*s* (Başaklar et al., 2019; Gembler et al., 2019; Gembler and Volosyak, 2019; Thielen et al., 2021; Martínez-Cagigal et al., 2023). This short selection time offers a user-friendly experience and further enhances the applicability of BCI in operational settings.

Indeed, ensuring a positive end-user experience is a critical concern when implementing reactive BCI systems. In this study, we aimed to address this concern by improving visual comfort, allowing users to engage with the system for extended periods. Unsurprisingly, subjective findings indicated that the *burst* codes were more comfortable than the m-sequence codes. This can be attributed to the fact that our *burst* codes, displayed at an approximate frequency of 3Hz, involved fewer flashes and emitted less luminance compared to the m-sequences. Additionally, reducing the amplitude depth led to higher visual comfort, as we decreased the intensity of flickers, consistent with previous studies (Ladouce et al., 2021, 2022). Interestingly, this reduction in amplitude depth did result in a slight decrease in accuracy, but it proved to be a favorable trade-off for the *burst* codes. The performance drop was minimal, reaching only 1.4% when using 6 blocks of data. Furthermore, even with 40% luminance and only three blocks of calibration data (26.4 s of training data), the *burst* code still achieved an accuracy of 90.7%. These minor reductions in classification performance should be considered with the substantial improvement in user comfort accompanying *burst* c-VEP amplitude depth reduction. These results highlight the practicality and robustness of the *burst* code, particularly when compared to the m-sequence codes. Notably, the reduction in amplitude depth exacts a more substantial toll on accuracy for the m-sequence, dropping to 77% at best and 66% with the same three blocks of data.

Two main explanations can account for the observed differences in classification performance. Firstly, and consistently with our hypotheses, the *burst* code successfully leverages the brain’s response, eliciting robust event-related responses. Electrophysiological analyses revealed that the *burst* codes are associated with prototypical visual P1 event-related responses that are prominent over occipital recording sites (over primary visual cortex areas). The amplitude elicited by the *burst* codes (approximately 4.4 *μ*V) was found to be twice as high as that of the m-sequence (approximately 2.2 *μ*V). These latter amplitudes for the m-sequence align with those reported by Thielen et al. (2015). Furthermore, the ITC evoked by the *burst* exhibited significantly more robust and distinguishable responses when compared to those elicited by the m-sequence. The m-sequence c-VEP evokes more complex and diffuse neural responses compared to the discrete VEP observed in response to the presentation of *burst* sequences. This difference may be attributed to the higher rate of visual stimulation provided by the m-sequences. The rapid succession of visual stimulus onsets can induce overlapping responses, resulting in non-linear interactions (Nguyen et al., 2019). Another hypothesis that could explain the lower amplitude and coherence of VEP responses observed in the sequence conditions is that the neural populations underlying the VEP response have a refractory period. The shorter inter-stimulation periods of m-sequences may not provide sufficient time for neurons to reset to synaptic excitability thresholds (Regan, 1989). The consistency of these larger amplitude responses to the *burst* codes facilitates the learning and training of the CNN, as it is presented with more reliable and predictable inputs. Secondly, the utilization of *burst* codes reduces the computation cost and training time for the CNN. The *burst* code not only reduce the classification problem to a 2-class one but also achieved a more balanced distribution of both binary classes, 1 and 0. In the case of the m-sequence, which entails a large number of flashes, the balance between classes is almost perfect due to the rapid succession of flashes whose mean duty cycle approximates 50%. On the other hand, *burst* stimulations involve fewer flashes with longer time intervals than m-sequences, resulting in a decreased relative number of instances for the on-state “1” than the off-state “0”. As such, the present results suggest that the state imbalance in the *burst* codes at least partially contributes to their superior performance classification-wise.

Collectively, these findings demonstrate the promising potential of the *burst* code sequences design to advance BCI technology. The *burst* codes sequences exhibit high classification accuracy, fast selection and training time while only requiring minimal training data. These findings emphasize the viability of the *burst* c-VEP design for practical applications that go beyond the laboratory. Furthermore, the reported differences in user experience related to the different types of codes demonstrate the critical importance of considering end-user satisfaction and highlight the advantages of utilizing *burst* codes with reduced amplitude depth to develop BCI solutions that will be effectively adopted and will present high retention rate of users in the long-term. The proposed *burst* c-VEP design strikes a favorable balance between performance and user experience. However, there are still areas that require further improvement, especially regarding its implementation. At present, *burst* codes elicit visually evoked potentials sparsely, at an interval of approximately 300ms. As a result, there are fewer flashes and therefore occurrences of the on-state ‘1’ in the *burst* code sequences than in the m-sequence codes. The reduced number of on-state instances offers certain advantages, notably in terms of improved user experience as both visual stimulation frequency and intensity are reduced. However, incorrect decoding of a VEP response associated with a flash (on-state ‘1’) is more punishing for the *burst* codes in contrast to the m-sequence codes that contain more instances of the on-state ‘1’. Indeed, the scarcity of flashes in *burst* code sequences may be detrimental to the reconstruction of the entire code sequence, either delaying or disrupting correct classification. Nonetheless, the electrophysiological analyses have revealed that the *burst* code sequences elicit strong VEP responses that can be reliably decoded. Although the exact nature of this increase in VEP response has to be further explored, an interesting hypothesis is that their scarcity may be an explanatory factor as it allows neural populations to complete their refractory period and therefore be more sensitive to following stimuli. In future works, the inter-flash interval of *burst* codes could be manipulated to find an optimal trade-off between performance and user experience. This adjustment would enable us to increase the flash rate from 3Hz to 5Hz, thus improving the robustness of the reconstruction and its discriminability. The *burst* generation algorithm can also be enhanced. So far, the strategy to generate such codes has been to create a pseudo-random batch of *n* codes and check their correlations pair-wise. This code generation process could be further improved through an iterative selection of *burst* codes that have a low correlation with an initial code, thus creating codes with good pairwise correlation by construction, rather by elimination of unfit codes from a codebook. Regarding the selection phase, our approach relies on a basic system where, if the same temporary prediction is outputted 60 times in a row, a closing prediction for the trial is triggered. To enhance the robustness and speed of this decision-making process, a more refined algorithm based on a Bayesian accumulation of Riemannian probabilities (Barthélemy et al., 2022) or a partially observable Markov decision process (Torre Tresols et al., 2023, 2022) could be employed. Such an approach would help improve the reliability of the decision phase while achieving faster outcomes. Another effective strategy to enhance user experience would be to minimize or even eliminate the need for training data altogether, thus bypassing the calibration phase. This could be achieved through transfer learning approaches as illustrated by (Thielen et al., 2021), who achieved high decoding accuracy despite minimal training data acquired from the user. By adopting transfer learning approaches, users interaction with a BCI system can be accelerated by using data from previous sessions or from other users for initializing or even fully pretraining the CNN. Although challenging, reducing calibration time is critical to provide a seamless experience of the BCI, and *burst* c-VEP as a reactive BCI paradigm, may be part of the solution to achieve this. Finally, the present work has used low-level visual contrast as an effective approach to elicit neural responses to state changes in primary visual cortical areas. However, future work may extend the design of *burst* c-VEP to other visual properties such as incorporating texture and motion (Ma et al., 2017), or even altering colors between states (Wei et al., 2016; Nezamfar et al., 2015) to further improve the signal-to-noise ratio of VEP responses and potentially improve user experience. Finally, the reliability and robustness of *burst* c-VEP make them ideal candidates to be put to the test of dry-EEG recordings outside of the laboratory (Dehais et al., 2020; Fairclough and Lotte, 2020).

## 5. ACKNOWLEDGMENT

This work was funded by AID (Powerbrain project), the AXA Research fund and ANITI (Chair for Neuroadaptive Technology).

## Footnotes

1 https://www.psychopy.org/

2 https://github.com/sccn/labstreaminglayer

3 https://github.com/neuroergoISAE/burst_codes

4 https://jasp-stats.org/

